# An unwanted association: the threat to papaya crops by a novel potexvirus in northwest Argentina

**DOI:** 10.1101/2022.09.09.507323

**Authors:** Dariel Cabrera Mederos, Humberto Debat, Carolina Torres, Orelvis Portal, Margarita Jaramillo Zapata, Verónica Trucco, Ceferino Flores, Claudio Ortiz, Alejandra Badaracco, Luis Acuña, Claudia Nome, Diego Quito-Ávila, Nicolás Bejerman, Onias Castellanos Collazo, Aminael Sánchez-Rodríguez, Fabián Giolitti

**Author notes:** **Authors for correspondence:** Aminael Sánchez-Rodríguez, Dariel Cabrera Mederos.

## Abstract

An emerging virus isolated from papaya (*Carica papaya*) crops in northwestern (NW) Argentina was sequenced and characterized using next-generation sequencing. The resulting genome is 6,667-nt long and encodes five open reading frames in an arrangement typical of other potexviruses. This virus appears to be a novel member within the genus *Potexvirus*. Blast analysis of *RNA-dependent RNA polymerase* (RdRp) and *coat protein* (CP) genes showed the highest amino acid sequence identity (67% and 71%, respectively) with pitaya virus X. Based on nucleotide sequence similarity and phylogenetic analysis, the name papaya virus X is proposed for this newly characterized potexvirus that was mechanically transmitted to papaya plants causing chlorotic patches and severe mosaic symptoms. RT-PCR based detection of papaya virus X (PapVX) revealed that it is widely present in papaya crops from NW Argentina. The prevalence of PapVX, which seems to be restricted to the NW region of Argentina, and the fact that it has only been detected in this region could be associated with a recent emergence or adaptation of this virus to papaya in NW Argentina.

## INTRODUCTION

Papaya (*Carica papaya* L.) is an important fruit crop widely cultivated in tropical and subtropical regions (Yeh et al., 2007). Papaya ranks fourth among tropical fruits produced worldwide, after banana, mango, and pineapple (FAO, 2022). The adaptability of this plant and the acceptance of its fruits provide considerable market advantages to this crop. In Argentina, the diversification of tropical fruit species is highly promoted with the aim of contributing to the development of local economies. Papaya trees are grown mainly in Jujuy, Salta, Corrientes, Chaco, Formosa and Misiones provinces, where the temperature is favorable for its cultivation.

Diseases caused by viruses are the main obstacle to papaya production worldwide (Tennant et al., 2007; Tripathi et al., 2008). The expansion of agriculture into tropical forests is an important driver of global climate change and biodiversity loss (Laurance et al., 2014). In this sense, research works have addressed the influence of agricultural activity on climate change and its impact on plant populations, vectors, and the dynamics of different viral pathosystems (Jones, 2009; Ghini et al., 2011). In the northern region of Argentina, temperature increase may favor the development of tropical crops, by reducing frost damage risk. Nevertheless, the rapid advance of extensive agricultural frontiers in subtropical regions can favor the development of emerging diseases (Jones, 2009).

In northwestern (NW) Argentina, papaya plants showing leaf mottling and mosaic symptoms, which differed from those previously reported for PRSV (Cabrera Mederos et al., 2016), were detected. In initial analyses of these samples, filamentous virus-like particles were identified and purified from symptomatic leaves. So far, only one potexvirus species, papaya mosaic virus (PapMV) (Sit et al., 1989; Kreuze et al., 2020), has been reported to naturally infect papaya plants worldwide. In addition, babaco mosaic virus (BabMV), a recently characterized potexvirus, was shown to infect papaya by means of mechanical inoculations (Alvarez-Quinto et al., 2017). With the development of next-generation sequencing (NGS) technology, plant virus discovery has increased enormously (Blawid et al., 2017) and several emerging papaya viral diseases have been detected in recent years (Medina-Salguero et al., 2019; Alcalá-Briseño et al., 2020; Mumo et al., 2020; Rumbou et al., 2021). In this study, we characterized the genome sequence of a novel potexvirus infecting papaya plants, according to evolutionary analyses and the demarcation criteria defined for *Potexvirus* genus (Kreuze et al., 2020). This virus represents a new member within this genus. The name papaya virus X (PapVX) is proposed for this new potexvirus. Symptoms observed in papaya plants were described, and the incidence and geographical distribution of this emergent virus were determined, showing local dispersion in papaya crops from NW Argentina.

## MATERIALS AND METHODS

### Sample Collections and Virus Maintenance

Papaya plants showing leaf mottling and mosaic were first observed in plantations in Yuto, Jujuy, in 2013. Afterward, papaya orchards in NW (Jujuy and Salta) and northeast (NE) (Chaco, Corrientes, Formosa, and Misiones) provinces of Argentina were surveyed from 2017 to 2021. Part of the collected symptomatic and asymptomatic leaf samples was stored at −80°C, and another part was lyophilized and stored at −20°C until use. For virus maintenance, leaf samples were ground in phosphate buffer 0.01 M Na_2_HPO_4_, pH 7+0.1% Na_2_SO_3_, and 600-mesh silicon carbide as abrasive, and the first three expanded leaves were mechanically inoculated onto five healthy papaya cv. ‘Maradol roja’ seedlings. In addition, five plants were mock-inoculated (control). The inoculated plants were maintained in an insect-free greenhouse for symptom development and further description (Hull, 2014).

### Electron Microscopy

Fresh symptomatic and asymptomatic leaf samples collected from both papaya field plants, and inoculated plants were analyzed by electron microscopy, using leaf-dip and inclusion in Spurr resin technique (Kitajima, 1997). Leaf-dips were stained with 2% uranyl acetate (Sigma-Aldrich, USA). Moreover, ultrathin sections were double-stained in 0.4% w/v lead citrate (Sigma Aldrich, USA) and 2% w/v uranyl acetate. Tissue preparations were examined with a JEM 1200 EX II electron microscope (JEOL, Japan) and photographed.

### Purification of Virus-Like Particles, Library Preparation and Next-Generation Sequencing

Leaf tissue from papaya plants showing leaf mottling and mosaic symptoms was used for purification of virus-like particles (Figure 1 A). Plants were collected in 2017 and preserved at −80°C until use. Plant material was processed using a metagenomics approach based on virion-associated nucleic acids (VANA) (Filloux et al., 2015). Concentration of virus-like particles was performed according to Lockhart (1986), with modifications. Leaves were ground and mixed in a blender with 1:4 (w/v) of cold 0.05 M Tris-citrate (pH 7.4) containing 0.5% Na_2_SO_3_ (w/v), 1% polyvinylpyrrolidone (PVP, mol wt 40,000) (w/v), and 1% Triton X-100 (w/v). The homogenate was squeezed through muslin and clarified by blending for 20 s; then 25% chloroform (v/v) was added, and centrifugation was performed at 10,000 g for 10 min. The aqueous supernatant was collected, and the virus-like particles were concentrated by centrifugation in a 30% sucrose gradient at 136,000 g at 4°C for 1 h, in a Beckman SW45 Ti rotor. The pellet was resuspended by stirring in 1 ml of 0.05 M Tris-citrate (pH 7.4) containing 0.5% Na_2_SO_3_ (w/v) at 4°C for 1 h.

**Figure 1.**
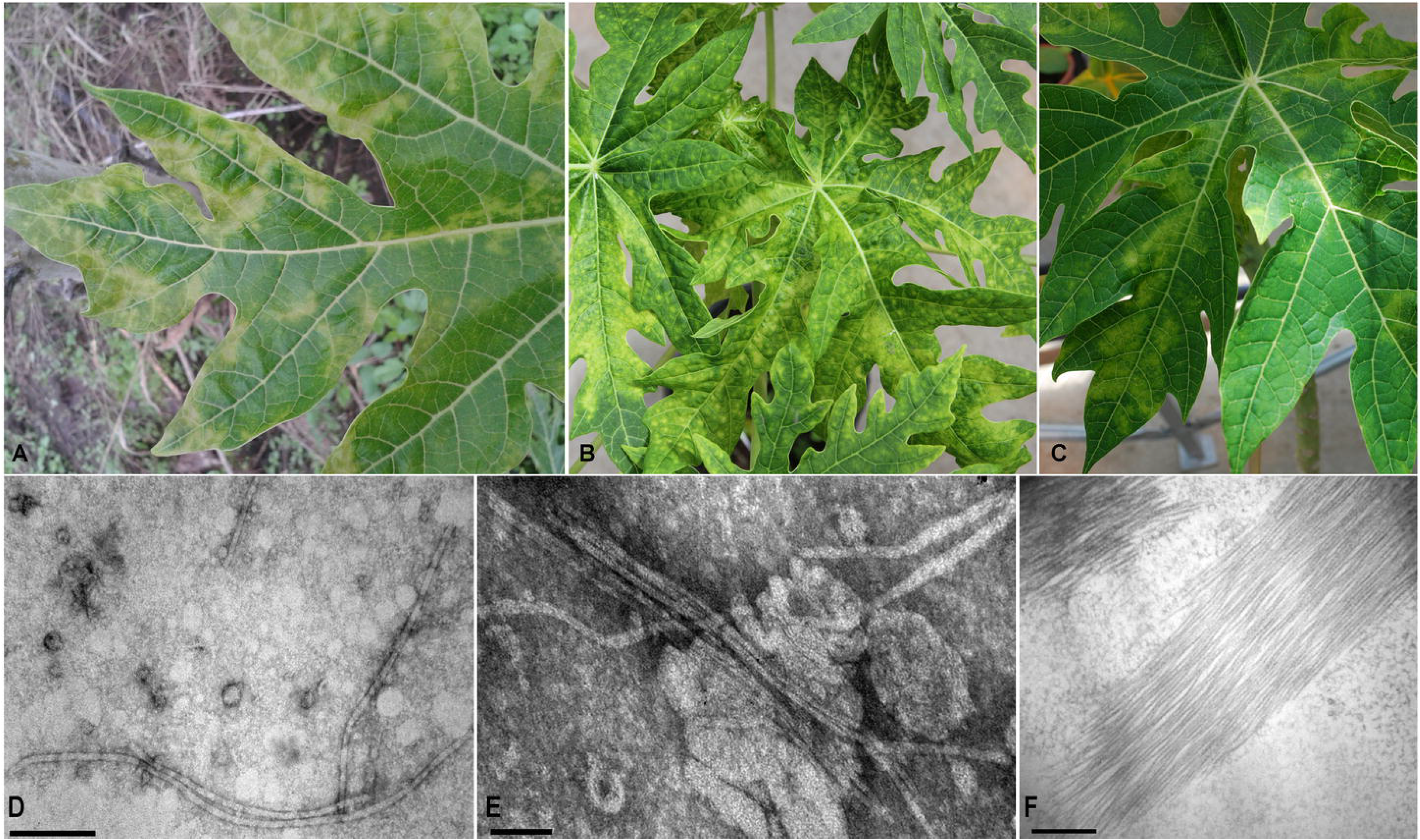
Papaya virus X in Argentina. (A) Symptoms (leaf mosaic, mottling, yellowing patches) induced in naturally and (B, C) mechanically infected papaya plants, respectively. (D, E) Electron micrographs of virus particles in leaf dip and (F) laminar aggregates in ultrathin sections.

For NGS, viral RNA was extracted from the suspension (200 μl) of concentrated particles using RNeasy Plant Mini Kit (Qiagen, Germany), according to the manufacturer’s protocol. RNA quality and concentration were determined by fluorometric quantitation Qubit 2.0 (Thermo Scientific, USA). RNA preparation (1 μg), subjected to ribosomal RNA depletion with Ribo-Zero rRNA Removal Kit (Illumina, USA), was used for library construction via TruSeq RNA Library Prep Kit v2 (Illumina, USA), according to the manufacturer’s protocol. The RNA was fragmented and copied into the first-strand cDNA using reverse transcriptase and random primers; then, the second-strand cDNA was synthesized using DNA polymerase I and RNase H. The cDNA was purified using Agencourt AMPure XP magnetic beads (Beckman Coulter, Inc., Indianapolis, IN, USA). A total of 3,084,077 paired-end 250-nt long reads were generated using the Illumina MiSeq platform in Castelar, Argentina.

### Sequence Processing and *de novo* Assembly

A trimming step was performed to remove adapters and low-quality reads using the Trimmomatic software version 0.39 (http://www.usadellab.org/cms/?page=trimmomatic) with standard parameters (Bolger et al., 2014). The processed reads were de novo assembled using Trinity software version 2.5.0 with standard parameters (Haas et al., 2013). The assembled contiguous sequences (contigs) were used as queries for BLASTX searching with standard parameters (https://blast.ncbi.nlm.nih.gov/Blast.cgi). The putative viral sequences obtained were curated by interactive reading mapping of filtered reads using the Bowtie 2 version 2.3.2 software (http://bowtie-bio.sourceforge.net/bowtie2/index.shtml) (Langmead and Salzberg, 2012). Genome organization, including identification of genomic features, was performed using the ORF finder available at National Center for Biotechnology Information (NCBI) and motif search tool from detected open reading frames (ORFs) (https://www.ncbi.nlm.nih.gov/orffinder/). Structural and functional domains of the detected gene products were predicted using the NCBI-CD-search tool available at https://www.ncbi.nlm.nih.gov/Structure/cdd/wrpsb.cgi, against the CDD v3.19 database with an expected value of 0.01.

### Phylogenetic Analysis

For phylogenetic analyses, sequences of *Alphaflexiviridae* members were obtained from the NCBI database (Table S1). Sequence alignments and comparisons were performed using MUSCLE (Edgar, 2004), via SeaView software version 5.04 (Gouy et al., 2010). To determine similarity among species, the selected sequences of RdRp were analysed using the circos method (Darzentas, 2000). Moreover, for species demarcation, pairwise identity analyses of gene sequences were performed using the MUSCLE-based pairwise alignment option, implemented in SDT software version 1.2 (Muhire et al., 2004). Phylogenetic analyses were performed on complete genome sequences and on *CP* and *RdRp* genes for both nucleotide (nt) and amino acid (aa) sequences. Sequence datasets were refined to conserve non-ambiguous sites using GBlocks with the least stringent conditions (Talavera and Castresana, 2007) in SeaView version 4 software. The evolutionary model for each dataset was estimated using the ModelFinder software (Kalyaanamoorthy et al., 2017), according to the Bayesian information criterion, selecting TVM+F+R6 (for complete genome), TVM+F+I+G4 (for CP nt), LG+G4 (for CP aa), GTR+F+R7 (for RdRp nt) and LG+F+R6 (for RdRp aa). The phylogenetic tree reconstruction was performed using maximum likelihood method implemented in the IQ-TREE version 1.6.12 software (Nguyen et al., 2015), available online (http://iqtree.cibiv.univie.ac.at/) (Trifinopoulos et al., 2016). The SH-like approximate likelihood ratio test (1000 replicates) (Guindon et al., 2010) and ultrafast bootstrap approximation (UFB) (10000 replicates) (Hoang et al., 2018) were used to evaluate the reliability of branches and groups obtained. In addition, a phylogenetic tree was also built using the RdRp partial sequences of isolates of the newly proposed papaya virus and other sequences of closely related species. The recombination detection program (RDP) 4 package (Martin et al., 2015) was used to detect possible recombinations across the whole-genome alignment of potexviruses using RDP, GENECONV, BOOTSCAN, MAXIMUM CHI SQUARE, CHIMAERA, SISCAN and 3SEQ methods, with default parameters.

### Host Range Assay

Mechanical inoculations were done by dusting carborundum (600-mesh) on test leaves followed by rubbing infected tissue homogenized in phosphate buffer as previously described (see Sample Collection and Virus Maintenance). Test plants included papaya, pitaya (*Hylocereus undatus*) Christmas cactus (*Schlumbergera truncata*), opuntia (*Opuntia* sp.), *Nicotiana glutinosa* and *Nicotiana rustica* (10 plants per species).

### Virus Detection

To determine the success of virus inoculation, each plant was RT-PCR tested at 60 days post-inoculation. Sense and antisense primers were designed based on the conserved aa motifs in the C-terminal region of the *RdRp* gene (ORF1) of the virus sequence obtained by Illumima MiSeq in this work and reference sequences of other related potexviruses, according to Van der Vlugt and Berendsen (2002). Designed primers were checked using the Primer-BLAST tool developed at NCBI (https://www.ncbi.nlm.nih.gov/tools/primer-blast/) and tested on the samples in which this virus was first detected by NGS.

Total RNA was extracted using Trizol method (Thermo Scientific, USA), according to the manufacturer’s instructions. The RNA was used for first-strand cDNA synthesis using Moloney murine leukemia virus (MMLV) reverse transcriptase enzyme (Promega, USA) and random primers (Biodynamics, Argentina). For virus detection, PCR reactions were performed using 4 μl of cDNA, 1.25 U of Go Taq G2 DNA polymerase enzyme (Promega, USA), 200 μM dNTPs and 0.3 μM of each primer in a final reaction volume of 50 μl. The PCR conditions included an initial incubation at 95°C for 2 min, followed by 35 cycles of denaturation at 95°C for 30 s, annealing at 55°C for 45 s, extension at 72°C for 60 s, and a final extension at 72°C for 10 min.

PCR products were separated on 1.0% agarose gel with TAE buffer (40 mM Tris base, 20 mM acetic acid, 1 mM EDTA pH 8.0). DNA bands were stained with Gel Red (Biotum Inc., USA), using GelDoc EQ (Bio-Rad Laboratories, USA) gel imaging system with the Quantity One analysis software version 4.5.1 (Bio-Rad Laboratories, USA) and scored by comparison to a 1kb DNA ladder (PBL, AR).

### Virus Distribution and Genetic Analyses

A virus survey was conducted to determine the distribution of this virus in the northern region of Argentina. Samples were collected from papaya orchard and family gardens (sampling sites) located in Jujuy, Salta, Chaco, Corrientes, Formosa and Misiones provinces. The geographic coordinates for sampling locations were obtained using a handheld global positioning system (GPS) (Table S2). Virus distribution was determined by molecular analysis of symptomatic and asymptomatic leaves previously collected and stored at −80°C. DIVA-GIS software version 7.5 was used to generate a map showing the virus distribution data. The virus was also tested in Ecuador, using papaya and babaco (*Vasconcellea x heilbornii*). A total of 120 papaya samples were collected from four commercial orchards distributed in three provinces, Santa Elena (n=30), Manabí (n=60), and Esmeraldas (n=30), whereas babaco samples were collected from a commercial plant nursery in Azuay province (n=30) and a commercial production operation in Pichincha province (n=30).

The RNA purification, cDNA synthesis and PCR conditions were the same as previously mentioned. The PCR products corresponding to symptomatic plants, obtained in duplicate, were purified using Wizard^®^ SV Gel and PCR Clean-Up System (Promega, USA), and then directly sequenced in both directions by Macrogen Inc. The obtained sequences were assembled with Staden package version 2.0.0b10 (https://staden.sourceforge.net), and submitted to the BlastN program (Altschul et al., 1990) for homology searching (http://blast.ncbi.nlm.nih.gov).

After sequence trimming, a 681-bp *RdRp* gene fragment was used to perform the phylogenetic analyses. The identity analyses among the obtained sequences were carried out using the MUSCLE-based pairwise alignment option implemented in SDT version 1.2 (Muhire et al., 2014). Multiple sequence alignments were used for ML phylogenetic reconstruction, as described above. The evolutionary model estimated was TIM3+F+I+G4, according to Bayesian Information Criterion. Papaya samples collected from Argentina were also tested, for two additional potexviruses *i.e*. PapMV and BabMV, using specific primers (Tuo et al., 2014; Alvarez-Quinto et al., 2017).

## RESULTS

### Papaya Symptoms and Particle Observations

Chlorotic patches and severe mosaic symptoms, similar to those observed in the field samples (Figure 1A), were observed in mechanically inoculated papaya plants (Fig 1 B, C). Initial symptoms were visible on young leaves at 15 days post-inoculation (dpi), and at 35 dpi the virus was able to systemically infect the inoculated plants. Filamentous virus-like particles were only observed in leaf dip preparation of symptomatic field samples as well as in mechanically inoculated plants (Figure 1 D, E). Particle aggregates were also observed in ultrathin sections (Figure 1 F).

### Sequence Assembly

A total of 2,221,356 reads with a mean length of 145 nt were left after trimming and quality control. The processed reads were assembled de novo using Trinity; 124,390 contigs were obtained, which were subsequently used as query for Blastx searches against the NCBI refseq virus proteins. A single 6,570-nt long contig obtained a significant hit (67.31% identity, E-value = 0.0) against the RNA-dependent RNA polymerase of pitaya virus X. Further inspection of the virus transcript and curation by iterative mapping of filtered reads resulted in a consolidated 6,667-nt virus sequence supported by 7,564 reads (mean size 108.8 nt), and a mean coverage 131.7X. The virus sequence showing features and identity with several members of the *Potexvirus* genus was further annotated.

### Genome Organization and Identification

The genome sequence of the detected papaya virus consisted of 6,667 nt, excluding a polyA tail, and is organized in five ORFs (Figure 2 A). Similar to other potexviruses, the ORF1 (nt 90–4,766) encodes the RdRp, consisting of 1558 aa with an estimated molecular weight of 177.6 kDa. The virus possesses a set of three partially overlapping ORFs: ORF2 (nt 4,766–5,455), ORF3 (nt 5,418–5,762) and ORF4 (nt 5,689–5,886), displaying the typical triple gene block (TGB) configuration of potexviruses. ORF5 (nt 5,890–6,561) encodes the putative coat protein (CP) (Table S3). The conserved aa motifs QDGAML (nt 4155–4172), HQQAKDE (nt 3588-3608) and TFDANTE (nt 4305–4325) were identified in the C-terminal region of the RdRp (van der Vlugt and Berendsen, 2002). Moreover, a catalytic RdRp domain, containing the characteristic core motif TG_X3_T_X3_NT_X22_GDD found in potexviruses (Martelli et al., 2007), was present near the C terminus of the RdRp. The conserved motifs, including viral methyltransferase (Vmethyltransf) (Rozanov et al., 1992), RNA helicase (Viral_helicase1) (Kreuze et al., 2020) and RdRp (Koonin, 1991), were also detected.

**Figure 2.**
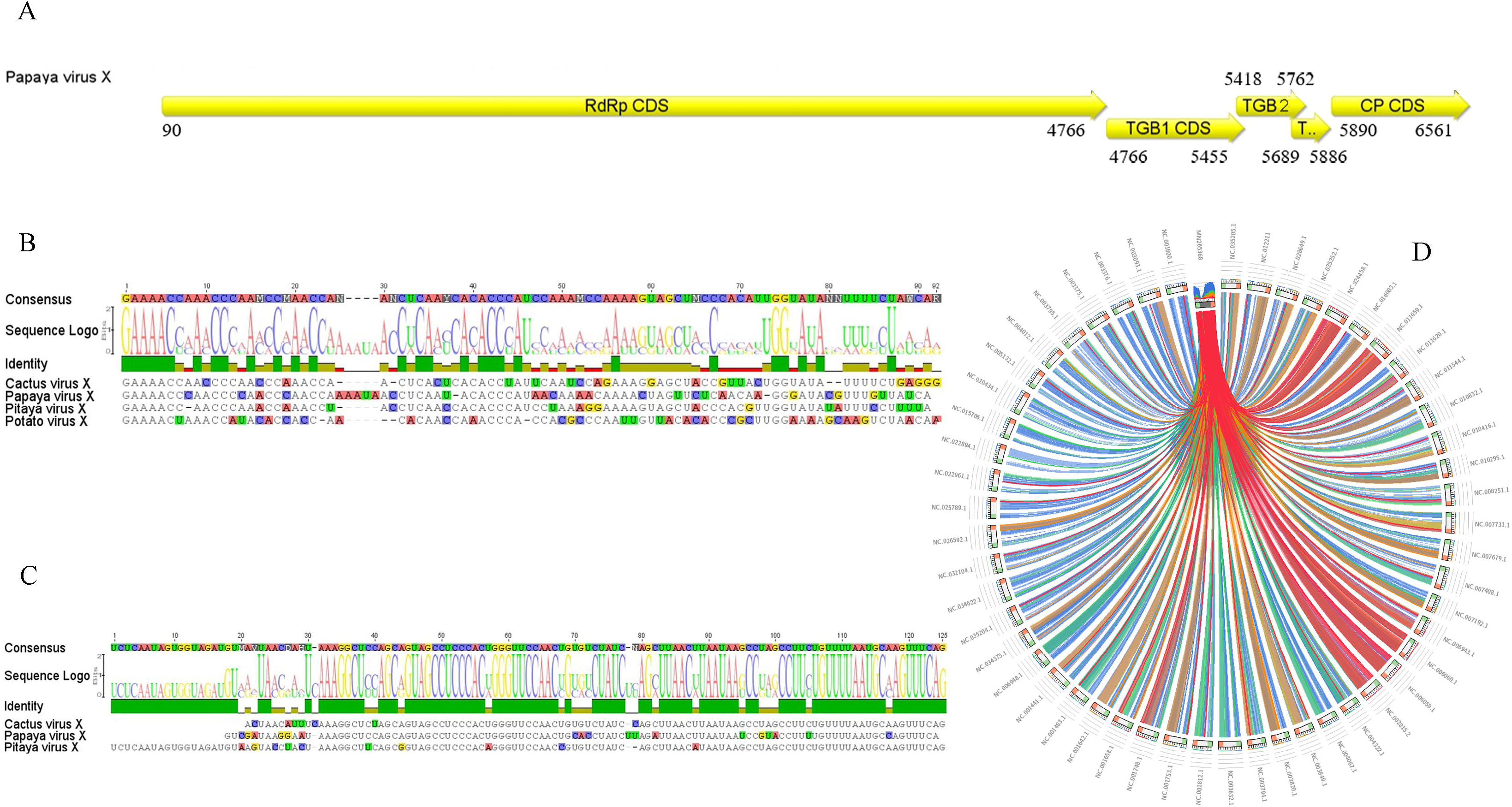
Schematic representation of the genome organization of papaya virus X. (A) The large open reading frames (ORF) are depicted by an open box (yellow boxes). The numbers indicate the nucleotide positions of the RNA segments for each mature protein in the polyprotein. (B, C) Sequence alignments between 5’ and 3’UTRs of PapVX and related potexviruses. The 3’UTR alignments are shown (excluding the polyA tail). (D) Sequence similarity plot obtained with Circos. Ribbons represent the local alignments produced by tblastx, their width the alignment length, and the colours the alignment percent similarity in four quartiles: blue<=58, green<=60, orange<=65, red>65.

Sequence alignments with potexviruses indicated the presence of untranslated regions (UTR) of 89 nt and 106 nt at the 5’ and 3’ ends, respectively. The hexanucleotide 5’-GAAAAC-3’, present at the 5’-UTR ends of most potexviruses, was observed (Figure 2 B). A putative polyadenylation signal, 5’-nt 6,572 AAUAAA-3’, important for the infectivity of potexviruses (Chen et al., 2005), was identified downstream of 3’-UTR (Figure 2 C). The presence of a typical 5’ end hexanucleotide at the 5’ UTR terminus and a putative polyA signal at the 3’ end indicates that the obtained virus sequence is coding-complete and tentatively corresponds to the complete genome of this novel virus. Similarity levels of the RdRp, expressed as Circoletto diagrams, confirmed a major relationship with those members of the *Potexvirus* genus (Fig 2 D). A BLAST search using RdRp and CP sequences revealed that the detected virus belongs to the *Potexvirus* genus. Pairwise sequence comparisons of the characterized virus and its closest relatives (pitaya virus X, zygocactus virus X, cactus virus X, schlumbergera virus X and opuntia virus X) revealed nt identities ranging from 66–68% for the RdRp and 64–67% for the CP. At the amino acids level, an average of 66.50% identity was observed for the RdRp and 66% for the CP (Table S4). In turn, the least conserved *TGB* genes showed nt identities ranging from 62 to 67% for TGB1, 56 to 58% for TGB2 and 55 to 56% for the TGB3 (data not shown). Based on the species demarcation criterion for potexviruses, which states that distinct species share <72% (nt) or <80% (aa) identity for the *RdRp* and *CP* genes (Adams et al., 2004; Kreuze et al., 2020), our results (Table S4) strongly support that the detected virus should be considered a new member within this genus, and the name papaya virus X is proposed (PapVX). The PapVX genome sequence was deposited in Genbank under accession number MN265368.1.

### Virus Phylogenetic Analysis

Maximum likelihood phylogenetic trees based on nt and aa sequences of *RdRp* from 61 alphaflexiviruses showed that PapVX clustered with potexvirus species pitaya virus X, zygocactus virus X, cactus virus X, schlumbergera virus X and opuntia virus X, in a highly supported group (Figure 3). The phylogenetic tree based on nt and aa sequences of the CP gene of potexviruses (45 species), and genome sequences (nt) (44 species) (Table S4) showed similar grouping among these viruses, with high support values in all trees (Figure 4, 5). No recombination events were detected among the genome sequences of potexvirus members analyzed.

**Figure 3.**
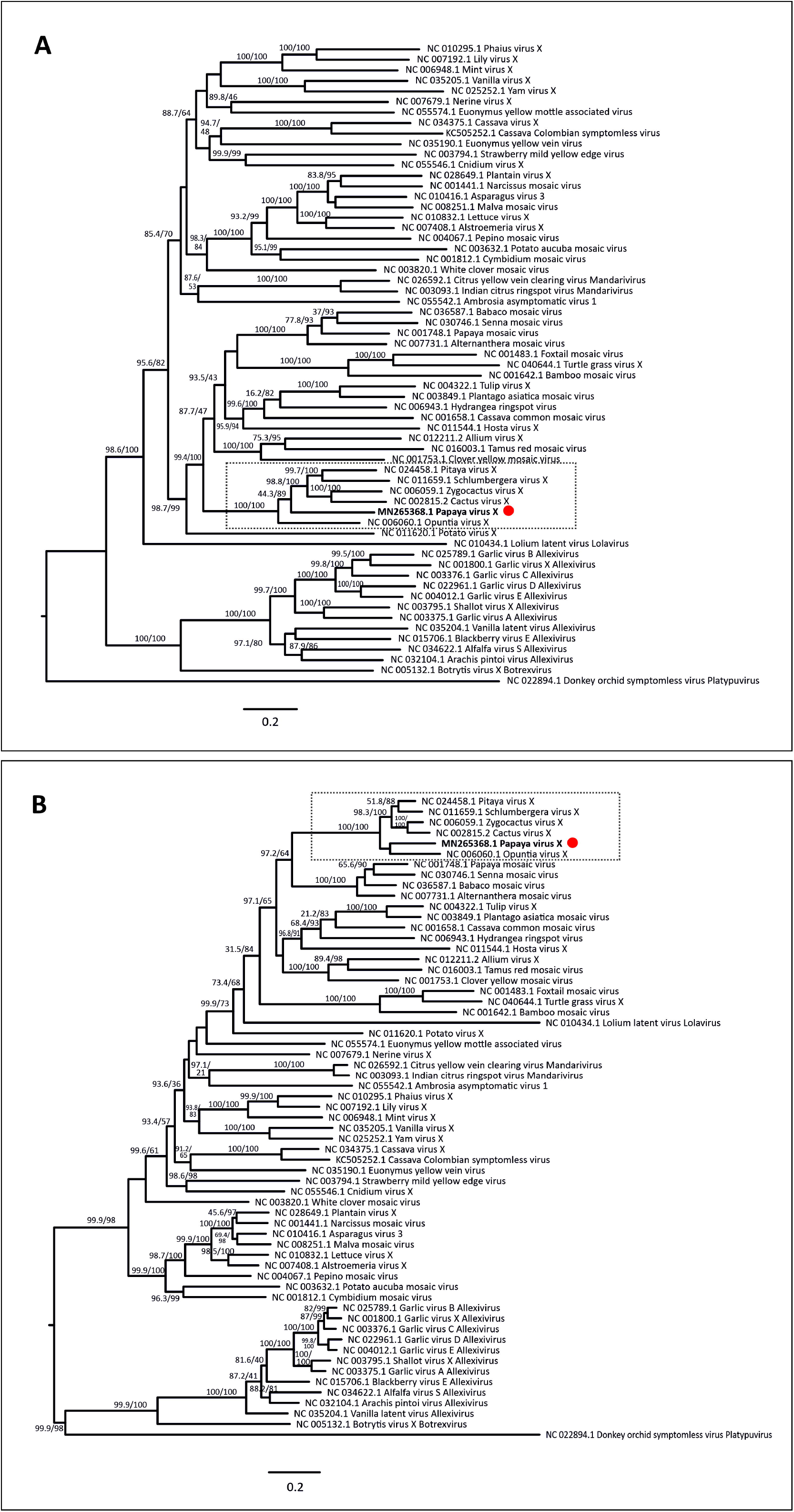
Phylogenetic tree of (A) nucleotide and (B) amino acid sequences of *replicase* gene of papaya virus X and virus species in the family *Alphaflexiviridae*. Branch lengths are proportional to genetic distances (nucleotide and amino acid substitutions per site). The SH-like / UFB support values are indicated when at least one of them is higher than 70%.

**Figure 4.**
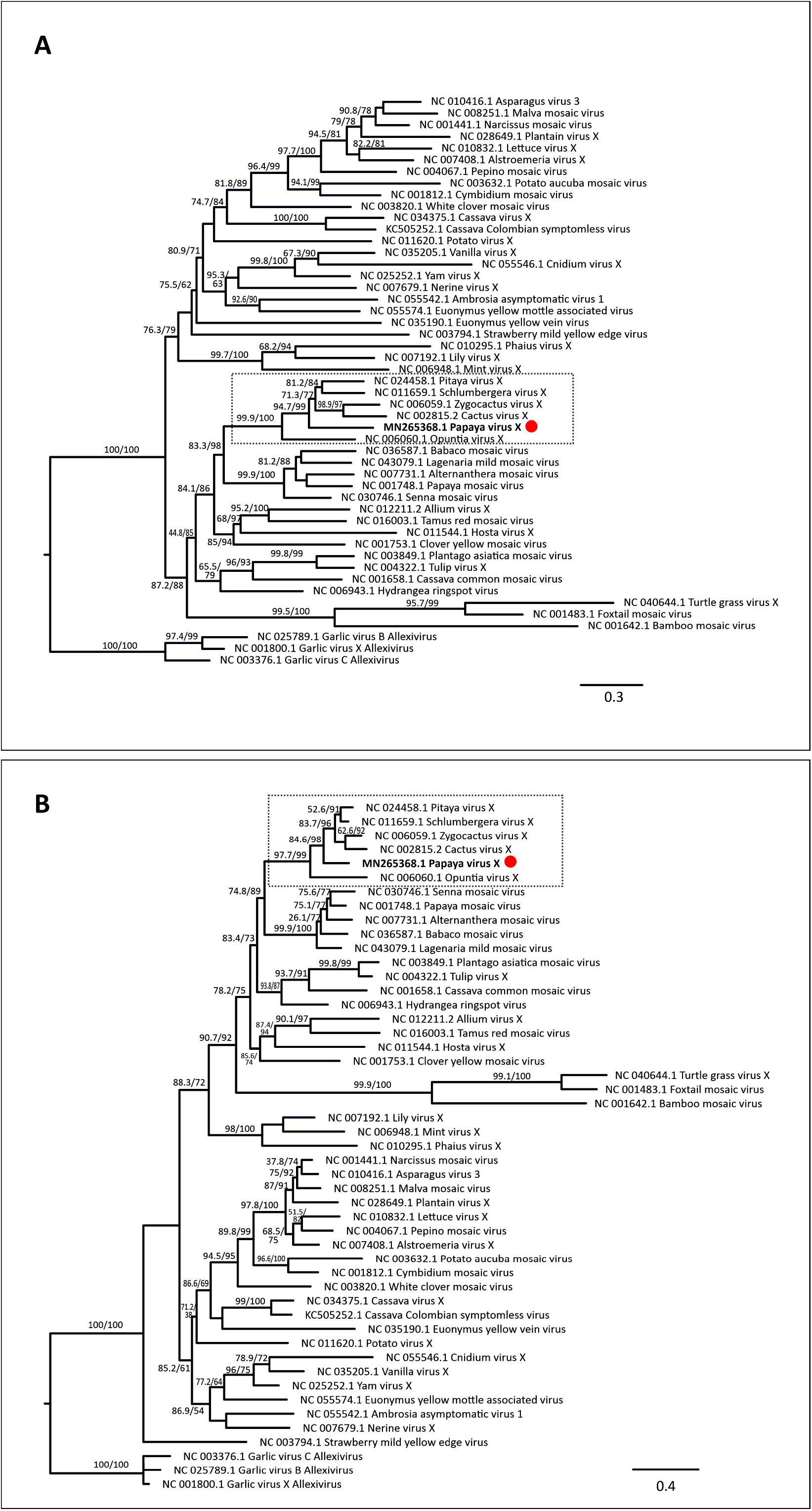
Phylogenetic tree of (A) nucleotide and (B) amino acid sequences of *coat protein* gene of papaya virus X and virus species in the genus *Potexvirus*. Branch lengths are proportional to genetic distances (nucleotide and amino acid substitutions per site). The SH-like / UFB support values are indicated when at least one of them is higher than 70%.

**Figure 5.**
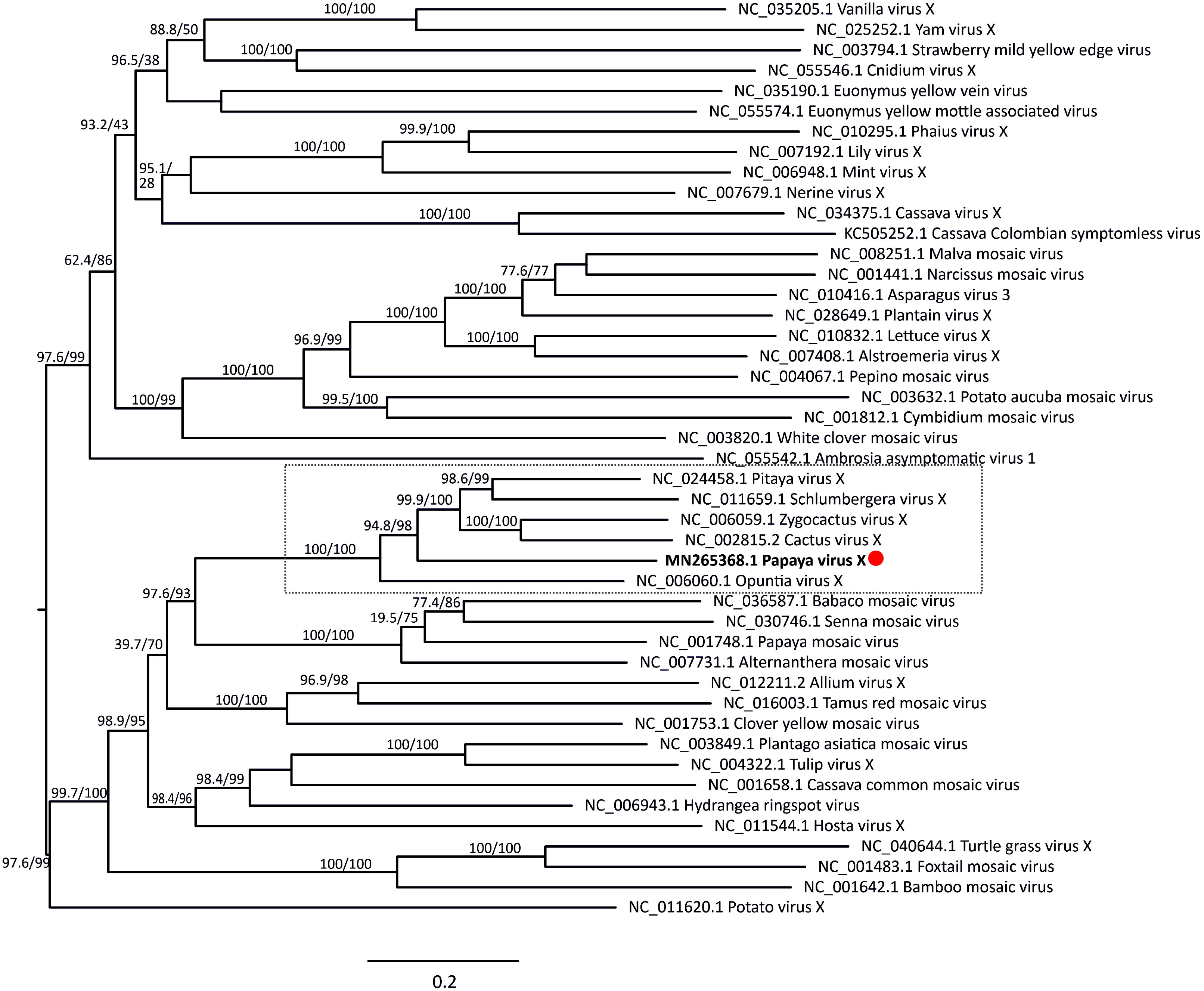
Phylogenetic tree of complete genome sequences of papaya virus X and virus species in the genus *Potexvirus*. Branch lengths are proportional to genetic distances (nucleotide substitutions per site). The SH-like / UFB support values are indicated when at least one of them is higher than 70%.

### Host Range Assay and Virus Detection

Back inoculation of PapVX to healthy papaya plants produced mosaic symptoms similar to those previously described. Virus symptoms were not observed in *H. undatus, S. truncate, Opuntia* sp., *N. glutinosa* and *N. rustica*. The presence of PapVX was molecularly confirmed only in papaya plants. The developed primer set (PapVX5 5’-CACCARCARGCNARRGATGA-3’ and PapVX1RC 5’-TCDGTGTTKG CRTCRAADGT-3’) resulted in amplification of ~737 bp PCR products from the terminal region of the *RdRp* gene. The inoculated papaya plants were also tested for PRSV (Cabrera Mederos et al., 2016), resulting negative for this virus infection.

### Virus Distribution and Genetic Analyses

PapVX was identified in 5 (23%) of the 21 localities sampled from the north of Argentina; only numerical tags were assigned to map locations where the virus was identified (Table S2; Figure 6). Of the 36 papaya plantations and family gardens (sampling sites) surveyed (18 from NE and 18 from NW regions), 8 sites were positive for PapVX, showing a prevalence of 22%. This virus was present exclusively in the NW region of Argentina. On the other hand, the potexviruses PapMV and BabMV were not detected in the inspected sites in Argentina. Moreover, PapVX was not detected in any of the papaya or babaco samples tested in Ecuador. PapVX detection primers and RT-PCR conditions were validated using BabMV-infected samples, which were successfully detected in babaco. The sequences obtained in this work were deposited in GenBank; details of the origins are shown in Table S2.

**Figure 6.**
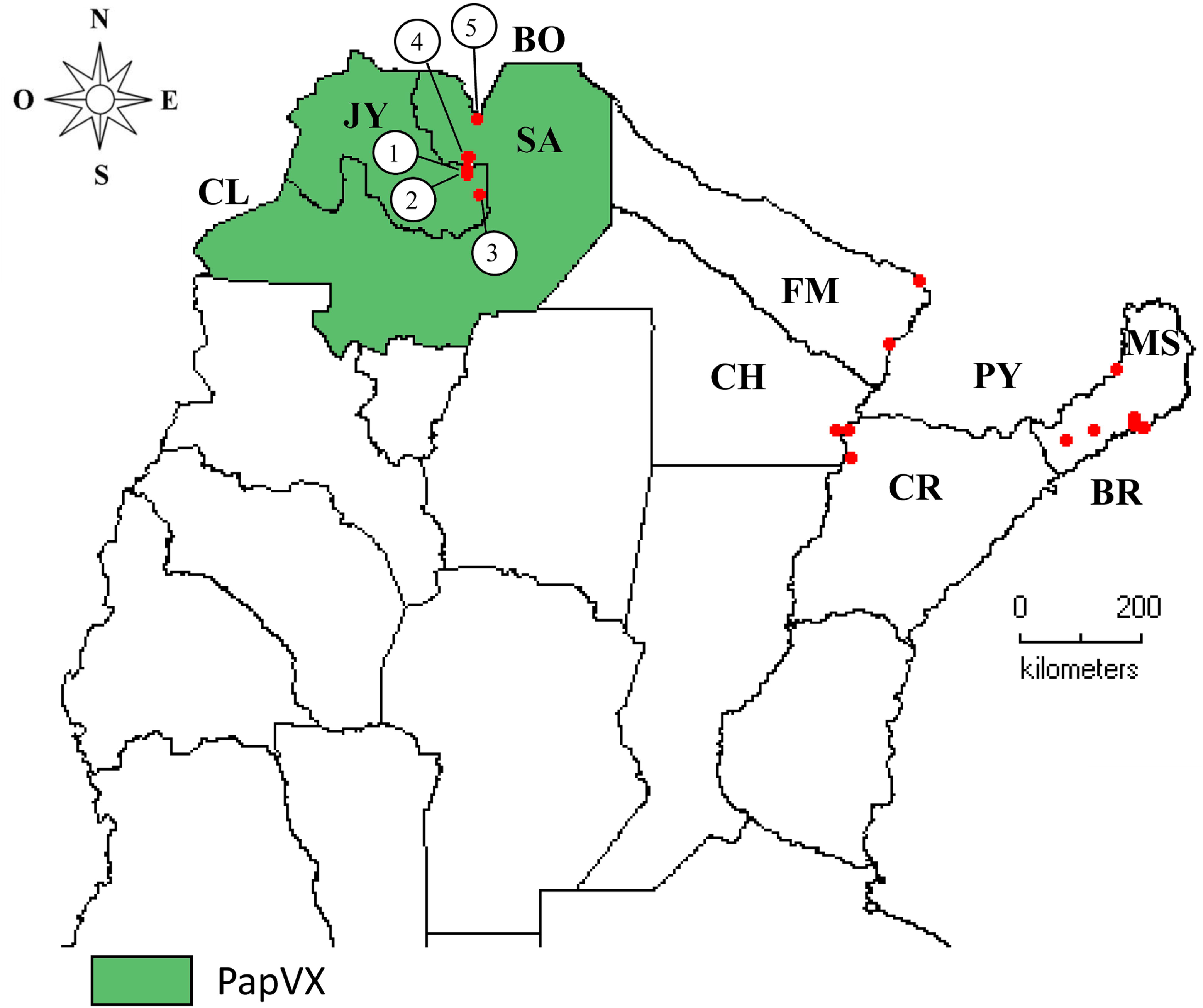
Geographical distribution of papaya virus X infecting papaya crops in Argentina. Locations where viral disease was detected are numbered from 1 to 5, and are shown in Suppl. Table 2.

The comparison between the partial *RdRp* gene sequences of PapVX obtained (GenBank accession numbers: ON007250, ON007251, ON007252, ON007253, ON007254) and the MN265368.1 showed nucleotide sequence identity values ranging from 82.99–99.71%, with the lowest identity values being detected between isolates from different locations. The pairwise analysis of the amino acid sequences showed values ranging from 98.56% to 100%. The highest similarities were observed between the sequences of isolates from nearby localities: isolates from Yuto (100%; ON007254 – MN265368.1) and Colonia Santa Rosa (100%; ON007251 – ON007252). In addition, in the phylogenetic tree of the partial *RdRp* gene sequences of PapVX, two subgroups were observed, one including sequences from Jujuy province (Yuto and Palma Sola) and the other from Salta province (Colonia Santa Rosa and Orán) (Figure S1).

## DISCUSION

The ICTV has officially recognized 45 species in the genus *Potexvirus* (https://talk.ictvonline.org/taxonomy/). In this work, we report the genetic and epidemiological studies of a novel potexvirus naturally infecting papaya crop in the NW region of Argentina. The sequencing strategy used in this study (TruSeq RNA Illumina protocol) targeted RNA viruses, whereas additional viral metagenomic analyses could help to understand the diversity of DNA viruses infecting papaya in the subtropical region of Argentina.

In this study, the symptoms observed in papaya plants were different from those attributed to PRSV infection, the only viral disease previously reported in this crop in Argentina (Cabrera Mederos et al., 2016). Several viruses may cause similar symptoms in one plant species and symptom expression in infected plants could be different according to the viral isolate or viral combinations (Chávez-Calvillo et al., 2016). In this sense, Mumo et al. (2020) reported similar symptoms in papaya plants to those attributed to PRSV infection (Tripathi et al., 2008), although PRSV was not detected. The biological methods for virus diagnosis are time-consuming, but they still hold very important in the characterization of plant viruses. In this regard, a clear association between PapVX presence and symptom expression was observed. Interestingly, PapVX was only reported in papaya plants with non-PRSV symptoms. The possible role of the potexvirus PapMV as a protective agent against PRSV and probably against other viruses was previously suggested (Cháves-Calvillo et al. 2016).

PapVX exhibits close genetic relationships and shares a common ancestor with pitaya virus X, zygocactus virus X, cactus virus X, schlumbergera virus X, opuntia virus X; for this reason, the name papaya virus X is proposed. Recombination and mutation events are major forces attributed to evolution in plant viruses and are associated with emergence of new variants and host adaptation (Fiallo-Olivé et al., 2019; Rubio et al., 2020); however, recombination events were not detected between PapVX and other potexviruses analyzed.

The presence of PapVX in papaya crops in different areas of NW Argentina suggests that the virus is extending its geographical distribution or that it has been present in papaya and/or other undetected host plants, which remains to be investigated. In this sense, the fact that this virus was detected exclusively in papaya plantations in Salta and Jujuy, but not in the NE region (Corrientes, Chaco, Formosa, and Misiones), suggests a recent emergence in this crop. Even with partial sequences, a geographical clustering was observed, indicating sustained transmission with still limited dispersion.

Although the phylogenetic relationships of PapVX with viruses isolated from cactaceous plants were confirmed, it was not experimentally transmitted to the plant species tested in this family (pitaya, Christmas cactus, and opuntia). In this sense, Alvarez-Quinto *et al*. (2017) showed that genetic diversity among potexviruses is not strictly related to host range. Although mechanical transmission could be a mode of virus dispersal, the successful infection of different plant species by mechanical transmission in the laboratory does not reflect what happens under field conditions. For example, babaco mosaic virus, a babaco-infecting potexvirus in Ecuador, does not infect papaya under field conditions, but mechanical transmission to papaya was successfully achieved (Alvarez-Quinto et al., 2017). Moreover, no insect vector has been reported to date in PapVX-related viruses.

The primer set PapVX5 and PapVX1RC resulted in the amplification of a 737-bp PCR products from the terminal region of the RdRp gene and will help to monitor the virus distribution and discover potential new hosts. These primers were employed to analyze papaya and babaco samples from Ecuador; results showed that all samples were negative for PapVX. The prospection of PapVX in papaya orchards from Bolivia, which borders the region where this virus was detected in Argentina, could be important to enhance the knowledge of PapVX dispersion. Linear patterns of papaya plants infected with PapVX were observed in the same crop row, which was potentially associated with mechanical labor (leaf removal) performed in the orchards, showing the importance of implementing cultural measures to avoid dispersion of PapVX within the crop. The potential vertical transmission of PapVX (via seeds), which plays a role in the long-distance dispersion of pathogens (Jones and Naidu, 2019), should be further characterized.

The use of high-throughput sequencing (HTS) to detect known and novel plant viruses or variants has extended considerably (Kutnjak et al., 2021), increasing our understanding of viral complexes and the diseases they cause. Known or emerging plant viruses are often reported in papaya, with more than 35 viral entities being known to cause disease in this crop worldwide. Although most of these viruses have a wide host range, such as TSWV and CMV, other viruses like PMeV and PapVQ, have only been detected in papaya (Sa Antunes et al., 2016; Cornejo-Franco et al., 2018), which could be similar to that observed for PapVX. To unravel these epidemiological aspects associated with the host range, the analysis of native plants associated with the papaya production should be included in the virus prospection.

The tropics and subtropics provide many examples of new encounter situations between viruses and wild or cultivated plants (Roossinck and García-Arenal, 2015; Jones and Naidu, 2019). For example, in NW Argentina, several emerging viral diseases have been reported (Vaghi Medina et al., 2018; Reyna et al., 2021). Papaya production in Argentina has increased significantly in the last years; as expected, virus-like diseases have also become conspicuous in the major production areas. The results obtained in this investigation encourage future studies considering the potential of native species as host of plant viruses to provide a complete panorama of virus emergence and evolution in the northern region of Argentina.

## CONCLUSSION

A virus naturally infecting papaya, tentatively named PapVX, is herewith reported for the first time. Given the rate at which papaya planting materials are being exchanged between farmers, it is likely that this virus, although currently restricted to NW of Argentina, could spread to other papaya-growing areas in the region. Additional surveys in other papaya crops areas, including native plants from hotspot sites and neighboring countries, should be carried out to determine the presence of this virus. Moreover, further epidemiological studies (host range, seed transmission) are needed to understand the risk that PapVX poses to papaya and other crops in the region.

## AUTHOR CONTRIBUTIONS

DCM, OP, AS-R, and FG: conceptualization, writing – original draft. HD: Genome assembly. HD, CT, AS-R, and DCM: data curation and formal analysis. DCM, MJ, CF, CO, AB, LA, DQ- A, and FG: sample collection. DCM, AS-R, and FG: funding acquisition. DCM, HD, CT, OP, CN, VT, AS-R; DQ-A; NB, and FG: writing – revision and editing. All authors contributed to the article and approved the submitted version.

## FUNDING

This work was partially funded by the INTA (project I090), FONCYT-ANPCyT (PICT 2020-02868), and Universidad Técnica Particular de Loja.

## ACKNOWLEDGMENTS

We thank the papaya farmers in the five Argentine provinces for allowing us to conduct sampling in their properties and Florencia Moreno (IPAVE-INTA, Argentina) for technical assistance. We also acknowledge the support of the Universidad Técnica Particular de Loja (Ecuador) for the Open Access Publication Fund of these results. The NGS was performed at the genomics unit of INTA Castelar (Buenos Aires, Argentina). The bioinformatic and evolutionary analyses were performed at the UFYMA and UBA, Argentina.

## Supplementary Materials

**Table S1**. List of viruses used for phylogenetic inferences.

**Table S2.** Geographical location of papaya collection sites in Argentina and collection dates.

**Table S3.** Characteristics of deduced proteins encoded by papaya virus X (PapVX) genome determined by predictive algorithms.

**Table S4.** Sequence identities of papaya virus X (MN265368.1) against reference sequences of reported species from *Potexvirus* genus.

**Figure S1.** Phylogenetic tree of nucleotide sequences of the replicase gene (partial) of papaya virus X and of the most closely related virus species in the genus *Potexvirus*. Branch lengths are proportional to genetic distances (nucleotide substitutions per site). The SH-like / UFB support values are indicated at nodes and tree is midpoint-rooted.

## Notes

### Competing Interest Statement

The authors have declared no competing interest.

