## Supplementary figures and images for "An unwanted association: the threat to papaya crops by a novel potexvirus in northwest Argentina"

### Figure S1

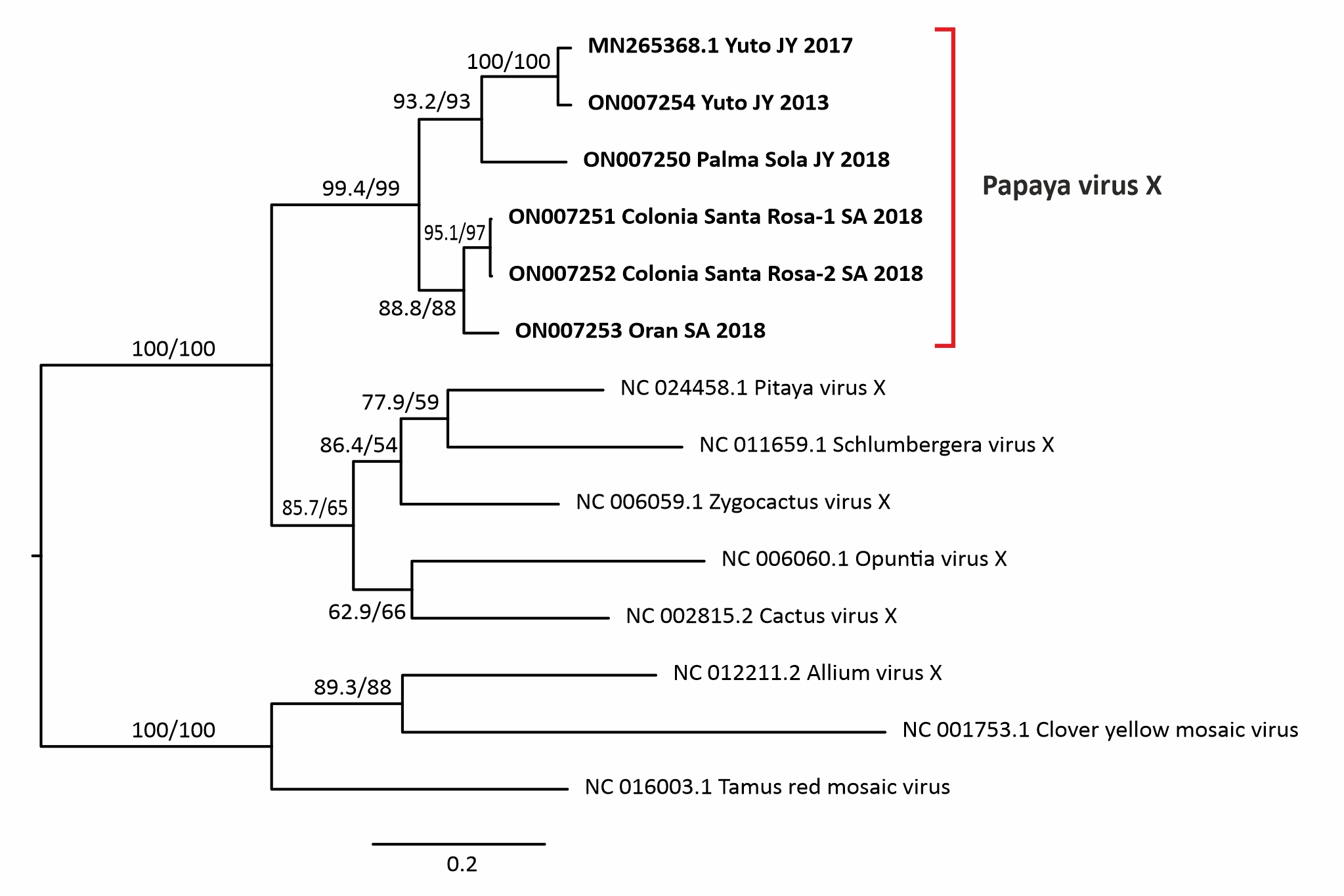
