## Supplementary material for "An unwanted association: the threat to papaya crops by a novel potexvirus in northwest Argentina": Table S1

**Table S1.** List of viruses used for phylogenetic inferences.

| **Species** | **NCBI Accession number** | **Genus** |
| --- | --- | --- |
| *Pitaya virus X* | NC_024458.1 | *Potexvirus* |
| *Zygocactus virus X* | NC_006059.1 | *Potexvirus* |
| *Cactus virus X* | NC_002815.2 | *Potexvirus* |
| *Schlumbergera virus X* | NC_011659.1 | *Potexvirus* |
| *Opuntia virus X* | NC_006060.1 | *Potexvirus* |
| *Allium virus X* | NC_012211.2 | *Potexvirus* |
| *Hosta virus X* | NC_011544.1 | *Potexvirus* |
| *Hydrangea ringspot virus* | NC_006943.1 | *Potexvirus* |
| *Tulip virus X* | NC_004322.1 | *Potexvirus* |
| *Plantago asiatica mosaic virus* | NC_003849.1 | *Potexvirus* |
| *Cassava common mosaic virus* | NC_001658.1 | *Potexvirus* |
| *Tamus red mosaic virus* | NC_016003.1 | *Potexvirus* |
| *Clover yellow mosaic virus* | NC_001753.1 | *Potexvirus* |
| *Alternanthera mosaic virus* | NC_007731.1 | *Potexvirus* |
| *Papaya mosaic virus* | NC_001748.1 | *Potexvirus* |
| *Babaco mosaic virus* | NC_036587.1 | *Potexvirus* |
| *Senna mosaic virus* | NC_030746.1 | *Potexvirus* |
| *Vanilla virus X* | NC_035205.1 | *Potexvirus* |
| *Yam virus X* | NC_025252.1 | *Potexvirus* |
| *Euonymus yellow vein virus* | NC_035190.1 | *Potexvirus* |
| *Phaius virus X* | NC_010295.1 | *Potexvirus* |
| *Lily virus X* | NC_007192.1 | *Potexvirus* |
| *Mint virus X* | NC_006948.1 | *Potexvirus* |
| *Cassava virus X* | NC_034375.1 | *Potexvirus* |
| *Potato virus X* | NC_011620.1 | *Potexvirus* |
| *Strawberry mild yellow edge virus* | NC_003794.1 | *Potexvirus* |
| *Plantain virus X* | NC_028649.1 | *Potexvirus* |
| *Malva mosaic virus* | NC_008251.1 | *Potexvirus* |
| *Narcissus mosaic virus* | NC_001441.1 | *Potexvirus* |
| *Asparagus virus 3* | NC_010416.1 | *Potexvirus* |
| *Lettuce virus X* | NC_010832.1 | *Potexvirus* |
| *Alstroemeria virus X* | NC_007408.1 | *Potexvirus* |
| *Cymbidium mosaic virus* | NC_001812.1 | *Potexvirus* |
| *Pepino mosaic virus* | NC_004067.1 | *Potexvirus* |
| *Potato aucuba mosaic virus* | NC_003632.1 | *Potexvirus* |
| *White clover mosaic virus* | NC_003820.1 | *Potexvirus* |
| *Nerine virus X* | NC_007679.1 | *Potexvirus* |
| *Euonymus yellow mottle associated virus* | NC_055574.1 | *Potexvirus* |
| *Cnidium virus X* | NC_055546.1 | *Potexvirus* |
| *Bamboo mosaic virus* | NC_001642.1 | *Potexvirus* |
| *Foxtail mosaic virus* | NC_001483.1 | *Potexvirus* |
| *Turtle grass virus X* | NC_040644.1 | *Potexvirus* |
| *Ambrosia asymptomatic virus 1* | NC_055542.1 | *Potexvirus* |
| *Cassava Colombian symptomless virus* | KC505252.1 | *Potexvirus* |
| *Lagenaria mild mosaic virus* | NC_043079.1 | *Potexvirus* |
| *Vanilla latent virus* | NC_035204.1 | *Allexivirus* |
| *Alfalfa virus S* | NC_034622.1 | *Allexivirus* |
| *Arachis pintoi virus* | NC_032104.1 | *Allexivirus* |
| *Garlic virus B* | NC_025789.1 | *Allexivirus* |
| *Garlic virus D* | NC_022961.1 | *Allexivirus* |
| *Blackberry virus E* | NC_015706.1 | *Allexivirus* |
| *Garlic virus X* | NC_001800.1 | *Allexivirus* |
| *Garlic virus E* | NC_004012.1 | *Allexivirus* |
| *Shallot virus X* | NC_003795.1 | *Allexivirus* |
| *Garlic virus A* | NC_003375.1 | *Allexivirus* |
| *Garlic virus C* | NC_003376. 1 | *Allexivirus* |
| *Indian citrus ringspot virus* | NC_003093.1 | *Mandarivirus* |
| *Citrus yellow vein clearing virus* | NC_026592.1 | *Mandarivirus* |
| *Lolium latent virus* | NC_010434.1 | *Lolavirus* |
| *Botrytis virus X* | NC_005132.1 | *Botrexvirus* |
| *Donkey orchid symptomless virus* | NC_022894.1 | *Platypuvirus* |
