## Supplementary material for "An unwanted association: the threat to papaya crops by a novel potexvirus in northwest Argentina": Table S2

**Table S2.** Geographical location of papaya collection sites in Argentina and collection dates.

| **Province** | **Department** | **Location** | **Date** | **GPS data** | **Map^a^** |
| --- | --- | --- | --- | --- | --- |
| Jujuy | Ledesma | Yuto | 2013 | -23.639104, -64.475260 | 1 |
| Jujuy | Ledesma | EEA INTA Yuto | 2017 | -23.586567, -64.507023 | 2 |
| Jujuy | Santa Bárbara | Palma Sola | 2018 | -23.970503, -64.294820 | 3 |
| Jujuy | Ledesma | Paraje Río Piedras | 2018 | -23.401801, -64,506167 |  |
| Salta | Orán | Colonia Santa Rosa | 2018 | -23.401801, -64.452116 | 4 |
| Salta | Orán | Peña Colorada | 2018 | -22.811360, -64.350422 | 5 |
|  |  |  | 2018 | -22.812280, -64.351546 |  |
| Chaco | San Fernando | Resistencia | 2017 | -27.468875, -58.974200 |  |
| Formosa | Formosa | Formosa | 2019 | -26.192428, -58.179419 |  |
| Formosa | Pilcomayo | Clorinda | 2013 | -25.251871, -57.728898 |  |
| Corrientes | Corrientes | Corrientes | 2013 | -27.475492, -58.777925 |  |
| Corrientes | Corrientes | Empedrado | 2021 | -27.903315, -58.737316 |  |
| Misiones | Leandro N. Alem | Cerro Azul | 2015 | -27.640641, -55.504605 |  |
| Misiones | Oberá | Oberá | 2016 | -27.491442, -55.114986 |  |
| Misiones | 25 de Mayo | Paraje El Palmital, Colonia Aurora | 2019 | -27.286206, -54.489858 |  |
| Misiones | 25 de Mayo | Paraje Cerro Grande, Colonia Aurora | 2019 | -27.407544, -54.489872 |  |
| Misiones | 25 de Mayo | Paraje Las Limas, Colonia Aurora | 2019 | -27.326556, -54.353164 |  |
| Misiones | 25 de Mayo | Paraje El Progreso, Colonia Aurora | 2019 | -27.390458, -54.454375 |  |
| Misiones | 25 de Mayo | Paraje Alicia Baja, Colonia Aurora | 2019 | -27.444689, -54.367283 |  |
| Misiones | 25 de Mayo | Paraje Alicia Baja, Colonia Aurora | 2019 | -27.429650, -54.357339 |  |
| Misiones | 25 de Mayo | Paraje Alicia Baja, Colonia Aurora | 2019 | -27.453594, -54.339841 |  |
| Misiones | Montecarlo | Montecarlo | 2019 | -26.566604, -54.759699 |  |

^a^ Map references used in Figure 6.
