## Supplementary material for "An unwanted association: the threat to papaya crops by a novel potexvirus in northwest Argentina": Table S3

**Table S3**. Characteristics of deduced proteins encoded by papaya virus X (PapVX) genome determined by predictive algorithms.

| **ORF No^a^** | **Gene** | **Calculated *Mr* (kDa)** | **Highest scoring virus protein/*E*-value/query coverage (Blast P)** |
| --- | --- | --- | --- |
| 1 | *RdRp* | 1503.182 | PiVX/0.0/100% |
| 2 | *TGB1* | 220.555 | PiVX/3e^-113^/97% |
| 3 | *TGB2* | 108.636 | CVX/6e^-34^/98% |
| 4 | *TGB3* | 62.273 | PVX/5e^-15^/96% |
| 5 | *CP* | 214.090 | SchVX/3e^-109^/100% |

^a^ ORF numbers are represented in the 5' to 3' direction of the viral genome and correspond to those in Figure 3A.
