## Supplementary material for "An unwanted association: the threat to papaya crops by a novel potexvirus in northwest Argentina": Table S4

**Table S4.** Sequence identities of papaya virus X (MN265368.1) against reference sequences of reported species from *Potexvirus* genus.

| **Species** | **Whole genome**  (nt) | **ORF1**  **RdRp**  (nt/aa) | **ORF5**  **CP**  (nt/aa) |
| --- | --- | --- | --- |
| *Pitaya virus X* | 68.01 | 67.94/67.85 | 67.57/69.51 |
| *Zygocactus virus X* | 67.66 | 68.41/67.30 | 65.57/66.37 |
| *Cactus virus X* | 66.65 | 67.02/65.04 | 64.98/66.36 |
| *Schlumbergera virus X* | 66.26 | 66.94/67.81 | 66.97/69.96 |
| *Opuntia virus X* | 66.19 | 66.88/64.50 | 64.55/58.11 |
| *Allium virus X* | 60.05 | 60.87/ 48.36 | 57.21/36.77 |
| *Hosta virus X* | 60.39 | 61.49/44.89 | 55.66/36.11 |
| *Hydrangea ringspot virus* | 61.42 | 63.19/52.55 | 57.66/49.55 |
| *Tulip virus X* | 61.19 | 62.02/50.71 | 58.88/42.93 |
| *Plantago asiatica mosaic virus* | 61.15 | 62.56/49.89 | 58.19/43.07 |
| *Cassava common mosaic virus* | 59.98 | 60.55/49.12 | 57.84/ 45.29 |
| *Tamus red mosaic virus* | 59.07 | 59.53/46.34 | 57.96/42.73 |
| *Clover yellow mosaic virus* | 60.58 | 60.93/48.13 | 58.33/43.54 |
| *Alternanthera mosaic virus* | 59.60 | 60.46/50.47 | 60.47/46.34 |
| *Papaya mosaic virus* | 60.32 | 61.34/51.55 | 59.54/44.13 |
| *Babaco mosaic virus* | 59.39 | 60.05/50.46 | 57.12/44.65 |
| *Senna mosaic virus* | 59.84 | 60.92/51.78 | 57.07/45.32 |
| *Vanilla virus X* | 59.23 | 59.86/44.78 | 54.24/36.57 |
| *Yam virus X* | 59.83 | 59.52/45.67 | 62.50/38.79 |
| *Euonymus yellow vein virus* | 59.75 | 59.38/44.08 | 56.97/31.53 |
| *Phaius virus X* | 59.83 | 60.59/48.25 | 57.57/37.86 |
| *Lily virus X* | 60.08 | 61.23/47.48 | 56.02/37.0 |
| *Mint virus X* | 60.69 | 60.69/47.19 | 57.07/33.79 |
| *Cassava virus X* | 60.54 | 60.54/47.21 | 59.32/33.18 |
| *Potato virus X* | 58.64 | 59.28/48.52 | 57.64/38.12 |
| *Strawberry mild yellow edge virus* | 58.41 | 59.48/48.15 | 56.45/32.57 |
| *Plantain virus X* | 58.50 | 59.21/41.79 | 57.03/28.05 |
| *Malva mosaic virus* | 58.50 | 58.64/42.50 | 60.71/30.77 |
| *Narcissus mosaic virus* | 58.43 | 59.11/41.66 | 57.28/29.22 |
| *Asparagus virus 3* | 59.40 | 59.97/43.95 | 56.18/30.59 |
| *Lettuce virus X* | 59.82 | 60.87/43.07 | 57.39/32.74 |
| *Alstroemeria virus X* | 58.40 | 58.67/44.53 | 58.68/32.58 |
| *Cymbidium mosaic virus* | 59.03 | 60.75/44.79 | 54.10/33.95 |
| *Pepino mosaic virus* | 58.34 | 59.04/43.59 | 59.62/32.42 |
| *Potato aucuba mosaic virus* | 58.31 | 58.63/41.13 | 56.76/34.58 |
| *White clover mosaic virus* | 59.66 | 60.63/50.51 | 58.26/36.70 |
| *Nerine virus X* | 58.65 | 59.69/45.20 | 57.62/33.94 |
| *Euonymus yellow mottle associated virus* | 60.33 | 60.37/47.82 | 60.55/35.16 |
| *Cnidium virus X* | 60.03 | 60.09/47.71 | 56.15/30.23 |
| *Bamboo mosaic virus* | 59.09 | 60.34/47.22 | 58.28/28.90 |
| *Foxtail mosaic virus* | 59.09 | 60.23/47.87 | 56.59/26.19 |
| *Turtle grass virus X* | 59.16 | 59.62/46.78 | 56.45/22.43 |
| *Ambrosia asymptomatic virus 1* | 59.26 | 59.46/45.19 | 57.24/38.53 |
| *Cassava Colombian symptomless virus* | 60.63 | 61.87/46.59 | 58.40/34.09 |
| *Lagenaria mild mosaic virus* | - | - | 57.26/47.09 |
| **Threshold^a^** | **-** | **72/80** | **72/80** |

^a^ The thresholds are as defined by Adams et al. (2004) and Kreuze et al. (2020). The thresholds currently accepted by the ICTV as criteria for species demarcation are indicated in bold. ORF: open reading frame; RdRp: RNA-dependent RNA polymerase; CP: coat protein.
